# Core genome MLST for epidemiological and evolutionary analyses of phytopathogenic *Xanthomonas citri*

**DOI:** 10.1101/2022.12.20.521341

**Authors:** R Ragupathy, K. A. Jolley, C. Zamuner, J. B. Jones, J. Redfern, F. Behlau, H. Ferreira, M.C. Enright

## Abstract

*Xanthomonas citri* subspecies *citri* (XCC) is the cause of bacterial citrus canker, responsible for major economic losses to the citrus industry that includes sweet orange, lime and grapefruit production in regions including South America, United States, China and Japan. Other *X. citri* subsp. and pathovars are responsible for diseases in crops such as soy bean, common bean, mango, pomegranate and cashew. Tracing the spread of *X. citri* disease has been performed using several different typing methods over the years but recent studies using genomic sequencing have been key to understanding evolutionary relationships within the species including fundamental differences between XCC pathotypes.

In this study we developed a core genome multilocus typing scheme (cgMLST) for *X. citri* based upon 250 genomes comprising multiple examples of *X. citri* subsp. *citri* pathotypes A, A* and A^w^, *X. citri* subsp. *malvacearum* and *X. citri* pathovars *aurantifolii, fuscans, glycines, malvacearum, mangiferaeindicae, viticola, vignicola* and single isolates of *X. citri* pathovars *dieffenbachiae* and *punicae*. This dataset included genomic sequencing of 100 novel XCC isolates. The cgMLST scheme, based upon 1618 core genes across 250 genomes, has been implemented at PubMLST (https://pubmlst.org/organisms/xanthomonas-citri/). Grapetree minimum-spanning, and iTOL Neighbour-joining phylogenies generated from cgMLST data resolved almost identical groupings of isolates to a core genome SNP - based neighbour joining phylogeny taking 4 min, 15 min and 36 h respectively. These resolved identical groupings of XCC pathotypes and *X. citri* subsp. and pathovars.

*X. citri* cgMLST should prove to be an increasingly valuable resource for the study of this key species of plant pathogenic bacteria. Users can submit genomic and associated metadata to compare with previously characterised isolates at PubMLST.org to allow rapid characterization of local, national and global epidemiology of these pathogens and examine evolutionary relationships.

## Introduction

Bacterial citrus canker has a major economic impact on the production of all commercial citrus crops including oranges, limes, tangerines, lemons and grapefruit. Three pathotypes of canker are recognised – A, B and C. Type A, caused by *Xanthomonas citri* subsp. *citri* (XCC), is the most widespread and economically damaging whereas types B and C, caused by *X. citri* pv. *aurantifolii* have much reduced virulence on sweet orange and have very limited geographical spread^(1)^. Type A(^2, 3^) pathotype has the broadest host range and infects most of the economically important citrus plants worldwide, particularly causing a major economic burden upon the South American and Californian orange industries (^2, 4^). Two variants of pathotype A have evolved:-A* that can cause canker on all citrus but with some isolates that can only infect key lime (*Citrus aurantifolia*), and A^w^ that only infects key lime and alemow (*C. macrophylla*)^(1)^.

*Xanthomonas citri* subsp. and pathovars other than XCC infect other important crop species including common bean (*X. citri* pv. *fuscans*), Mexican lime (pv. *aurantifolii*), mango (pv. *mangiferaeindicae*), grape (pv. *viticola*), cotton (subsp. *malvacearum*), soybean (pv. *glycines*), Araceae (pv. *dieffenbachiae*), cashew (pv. *anacardii*), and pomegranate (pv. *punicae*). Previous genome sequencing studies have examined the evolution of *X. citri* pathovars and subsp.^(5)^ and subsp. *citri* pathotypes^(6)^ and these have produced robust phylogenies that clearly resolve clades corresponding to individual *X. citri* pathovars and XCC pathotypes. Genomic sequencing has also proven useful in investigations of host-pathogen interactions through identification of host-specific virulence factors^(7)^.

Whole genome sequencing has greatly advanced the study of the epidemiology and evolution of pathogenic bacteria, greatly improving upon the discriminatory power and portability of other approaches such as ribotyping or pulsed-field gel electrophoresis^(8)^. Genomic sequencing and analysis tools, developed primarily for the study of human bacterial pathogens to track and investigate outbreaks of disease caused by particularly virulent or antibiotic-resistant clones, can also be usefully employed for the study of bacterial plant disease epidemiology and evolution.

Whole genome sequences from isolates of pathogenic bacteria are usually compared using SNP (single nucleotide polymorphisms) based approaches that involve whole genome alignments. Such SNP based approaches have been used in recent studies of *Xanthomonas citri* biology(^9, 10^), however they involve identifying genomes of isolates from the literature, downloading their sequences followed by the computationally intensive alignment of multiple genomes to generate SNP profiles that are then used to produce phylogenetic trees using methods such as neighbour-joining, maximum parsimony or maximum likelihood. Core-genome multilocus sequence typing (cgMLST) uses whole genome sequence data to examine genetic similarities between isolates. It is based upon allelic variations at a large number of core genome loci that are present in all, or nearly all, members of a species^(11)^. It differs from other whole genome sequencing approaches in that it does not include non-core, accessory genes in comparing genomes and, as it examines variation in allelic profiles rather than core genome SNPs. In addition, cgMLST is computationally efficient, scalable, and suited to the representation of very large numbers of genomic comparisons. cgMLST schemes have been established for diverse range of human pathogens and some schemes contain many thousands of genomes. For example, the curated, open-source database PubMLST (https://pubmlst.org/) contains genomic and metadata from 655,340 genomes of >100 bacterial species and that of Enterobase (https://enterobase.warwick.ac.uk) contains 379,370 Salmonella and 237,066 *E. coli* / Shigella genomes and corresponding metadata alone (as of 23^rd^ November 2022).

In this study we describe a core-genome multilocus sequence typing (cgMLST) scheme and website resource that can be used to rapidly and easily identify *X. citri* subsp *citri* variants from genome sequences without the need for computationally- and time– consuming core-genome SNP extraction, genome alignment and phylogenetic comparisons. The *X. citri* cgMLST database at https://pubmlst.org/organisms/xanthomonas-citri represents an invaluable resource for tracking the spread of pathovars of this devastating pathogen that should also prove to be a useful, scalable, tool in future national and international citrus canker and other crop disease control efforts.

## Methods

### Bacterial isolates

101 *X. citri subsp. citri* isolates were obtained from Fundecitrus, Araraquara, Sao Paulo, Brazil, an association maintained by citrus growers and juice manufacturers from the state of São Paulo to conduct research, education and implementation of citrus crop protection. Isolate 306, corresponding to the previously sequenced genome of strain 306^(12)^ was re-sequenced as part of this study resulting in the sequencing of 100 novel isolates. These were sampled from citrus plants from 15 different countries and included 75 isolates from Brazil, four from South Korea, three each from Argentina and the USA, two each from China, New Zealand, and Paraguay, and one each from Australia, Fiji, France, India, Iran, Mauritius, Taiwan, Thailand and Uruguay. One isolate’s country of origin is unknown. Details of isolates are shown in Table 1. These isolates were all pathotype A isolates from sweet orange, with the exception of two pathotype A* isolates from key lime. Study bacteria were isolated between the years 1979 and 2015. Data on year of isolation were not available for 31 of the 100 isolates.

**Table 1.**
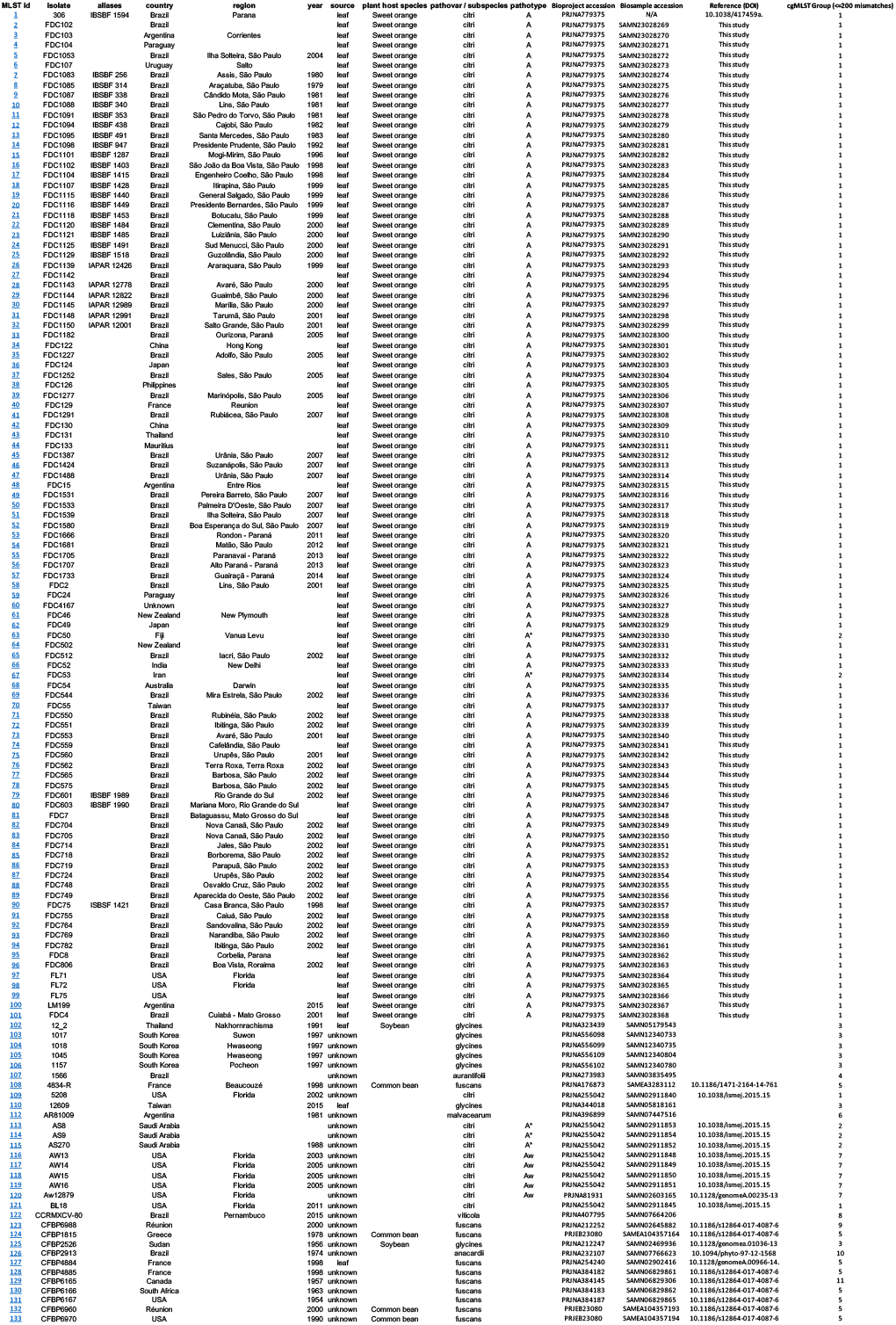
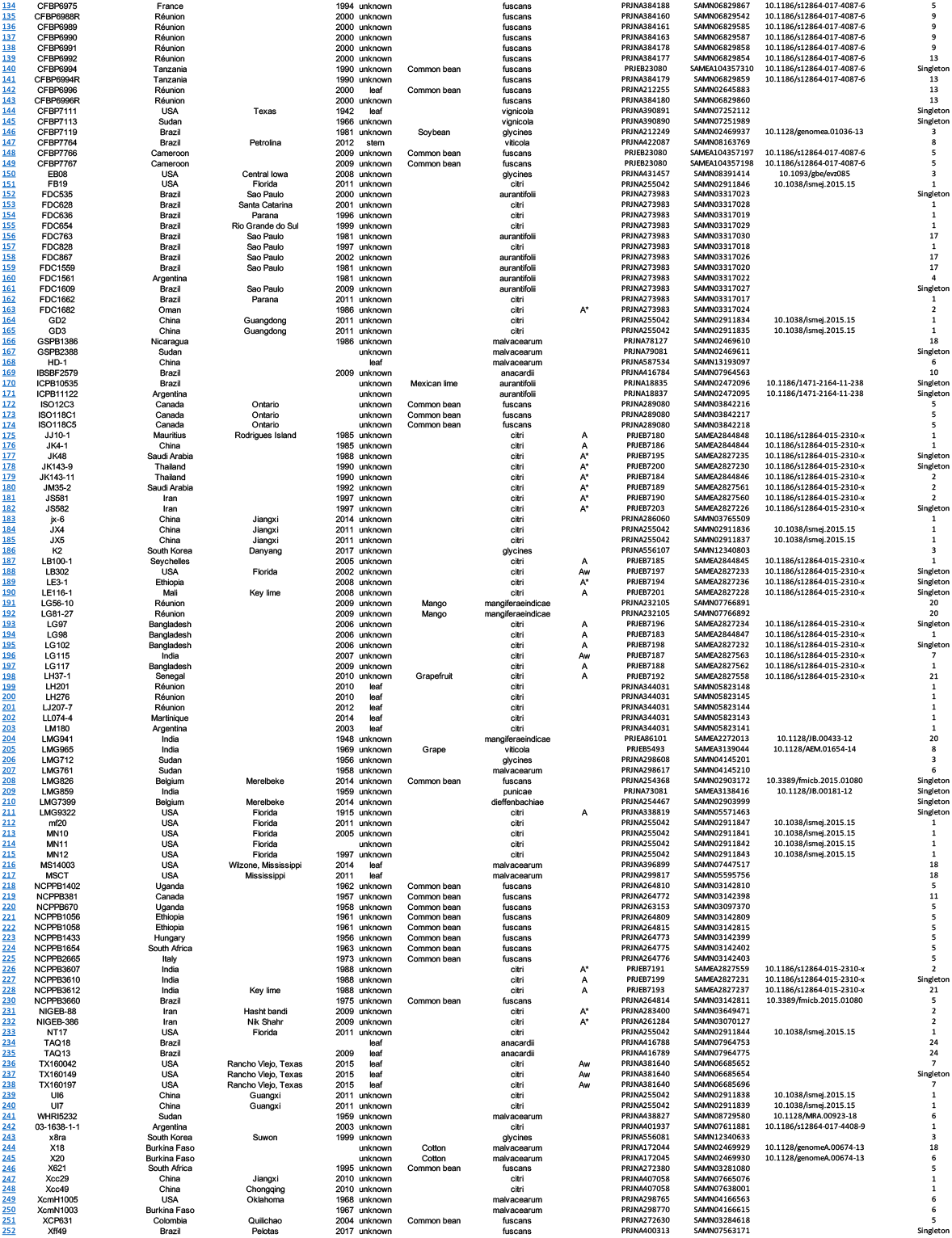
Details of isolates and genomes used in this study.

### Genomic DNA sequencing

Genomic sequencing (Illumina) was performed by MicrobesNG (University of Birmingham) from pure culture material stabilised in DNA / RNA Shield buffer (Zymo Research, CA, USA). A further 150 genome sequences, downloaded from the European Nucleotide Archive (ENA), were included for analysis from 65 XCC (comprising 12 pathotype A, 14 A* and 10 A^w^ isolates according to their cited literature source) and 9 *X. citri* pv. *aurantifolii*, 37 pv. *fuscans*, 12 pv. *glycines*, 12 subsp. *malvacearum*, 3 pv. *mangiferaeindicae*, 3 pv. *viticola*, 2 pv. *vignicola*, and 1 each of pv. *dieffenbachiae* and pv. *punicae*. Details on study isolates and those whose genomes were included in this study are shown in Table 1.

### Core genome MLST

Complete coding sequences were identified in the finished genome assembly of strain 306^(12)^ using Prokka^(13)^ with default settings. These were used in Roary^(14)^ to identify 1618 genes found in all 250 genomes. A BIGSdb database for *X. citri* was set up on the PubMLST website^(15)^ with loci defined for each of the identified core genes and named using a XCIT prefix and five digit identifier, ranging from XCIT00001 to XCIT01618. The database was seeded with the coding sequence found in strain 306 for each of these loci defined as allele 1. Allelic variants found in the 100 isolate locally sequenced dataset were then identified using the BIGSdb allele caller with thresholds of 98% identity over 98% alignment length compared to reference alleles. A further round of allele calling using the same parameters and all previously identified alleles as references was performed, followed by manual scanning to identify more variable alleles containing small indels. The database was then expanded to include all 250 isolates and alleles identified as above. Start codon positions were adjusted in nine loci as the codon identified in the reference genome was not found consistently across the dataset, whereas as an alternate consensus start codon was identified nearby. Core genome sequence types (cgSTs) were defined automatically by BIGSdb for profiles with fewer than 50 missing loci. Single-linkage cluster schemes were set up within the database to identify related isolates using a range of locus mismatch thresholds (200, 100, 50, 25, 10, and 5 locus mismatches).

### Phylogenetic trees

In common with previous studies of *X. citri* evolution and epidemiology we generated phylogenies based upon core-genome SNPs using a reference genome. We used the finished genome of *X. citri* pv. *citri* strain 306^(12)^ as a reference and kSNP3 v3.12^(16)^ to generate fasta nucleotide files of SNPs that were used in MEGA 11^(17)^ to generate NJ trees. Genomic DNA sequence data and associated metadata can be analysed using a variety of methods implemented at the Pubmlst website. Here we generated minimum-spanning trees from allelic profiles using GrapeTree^(18)^. Neighbour-joining trees based on concatenated nucleotides of cgMLST loci were generated using Interactive Tree of Life (iTOL)^(19)^. Both GrapeTree and iTOL plugins are implemented on the PubMLST website (pubmlst.org).

## Results

### cgMLST

1618 core genes (present in >99% isolates) were found among 250 *X. citri* isolate genomes. These genes were numbered XCIT00001 – XCIT01618. Allele calling of the initial 100 subset of records (from study isolates) resulted in isolates having between 99.4% and 100% of their loci with alleles designated. Core genome MLST (cgMLST) groupings of the 250 genomes uploaded to the PubMLST website were made based upon the number of allelic mismatches. This resulted in 171 groups of genomes with 5 or fewer mismatches (isolates tagged as Xc_cgc_5 at the PubMLST website), 113 with 10 or fewer (Xc_cgc_10), 53 with 50 or fewer (Xc_cgc_50), 39 with 100 or fewer (Xc_cgc_100) and 25 with 200 or fewer mismatches (Xc_cgc_200).

### cgMLST Groupings

The groupings of 250 isolates / genomes with fewer than 200 mismatches are shown in Table 1. Group 1 contained 132 subsp. *citri* isolates comprising 104 pathotype A isolates (pathotype data is missing for 28 isolates in this group); group 2 – 12 subsp. *citri* genomes, all of isolates with pathotype A*; group 3 comprised 12 pv. *glycines*; group 4 - two pv. *aurantifolii*; group 5 - 24 *pv. fuscans*; group 6 - seven subsp. *malvacearum*, group 7 – eight pv. *citri* isolates with pathotype A^w^; group 8 – three pv. *viticola* isolates; group 9 – five *pv. fuscans* isolates; group 10 – two pv. *anacardia* isolates; group 11 – two pv. *fuscans* isolates; group 13 – four pv. *fuscans* isolates; group 17 – three pv. *aurantifolii* isolates; group 18 – four subsp. *malvacearum* isolates; group 20 – three pv. *mangiferaeindicae* isolates; group 21 – two pv. *citri* isolates of unknown pathotype; and group 24 – two pv. *anacardia* isolates. 23 isolates had no close matches using any of the allelic mismatches groupings above and these are referred to as singleton isolate genomes in Table 1. Other cgMLST groupings (and all other genomic and metadata) can be found by selecting the hyperlink for each isolate in the ‘MLST id’ column of Table 1.

### X. citri phylogeny

A neighbour-joining tree of concatenated MLST allelic sequences of the 250 Xanthomonas isolates is shown in Figure 1. This phylogeny was generated using the iTOL plugin at pubmlst.org. It clearly distinguishes between individual *X. citri* pathovars (coloured) with genomes of isolates belonging to the same pathovar grouping together although some sub-groupings are evident. This is most marked for *X. citri* pv. *fuscans* isolate genomes that are represented by three clades that include isolates from a 2008 study by Alavi *et* al ^(20)^ and a 2015 study by Aritua *et al*^(21)^. These correspond to isolates from three lineages originally named pv. phaseoli and pv. phaseoli GL 1 and pv. phaseoli GL fuscans GL2 and 3. Genomes from isolates of pathovar *aurantifolii* resolve as two main clades with the smaller group sharing more genetic similarity to members of the *anacardii* group than to members of the other *aurantifolii*. Overall, this phylogeny displays a high degree of congruence with a neighbour-joining phylogeny based on core genome SNPs with the same groupings of genomes and only superficial differences in tree structure (Suppl. Figure 1). A minimum-spanning tree generated using Grapetree based upon cgMLST allele data shows identical groupings to those made using both methods (Figure 2).

**Figure 1.**
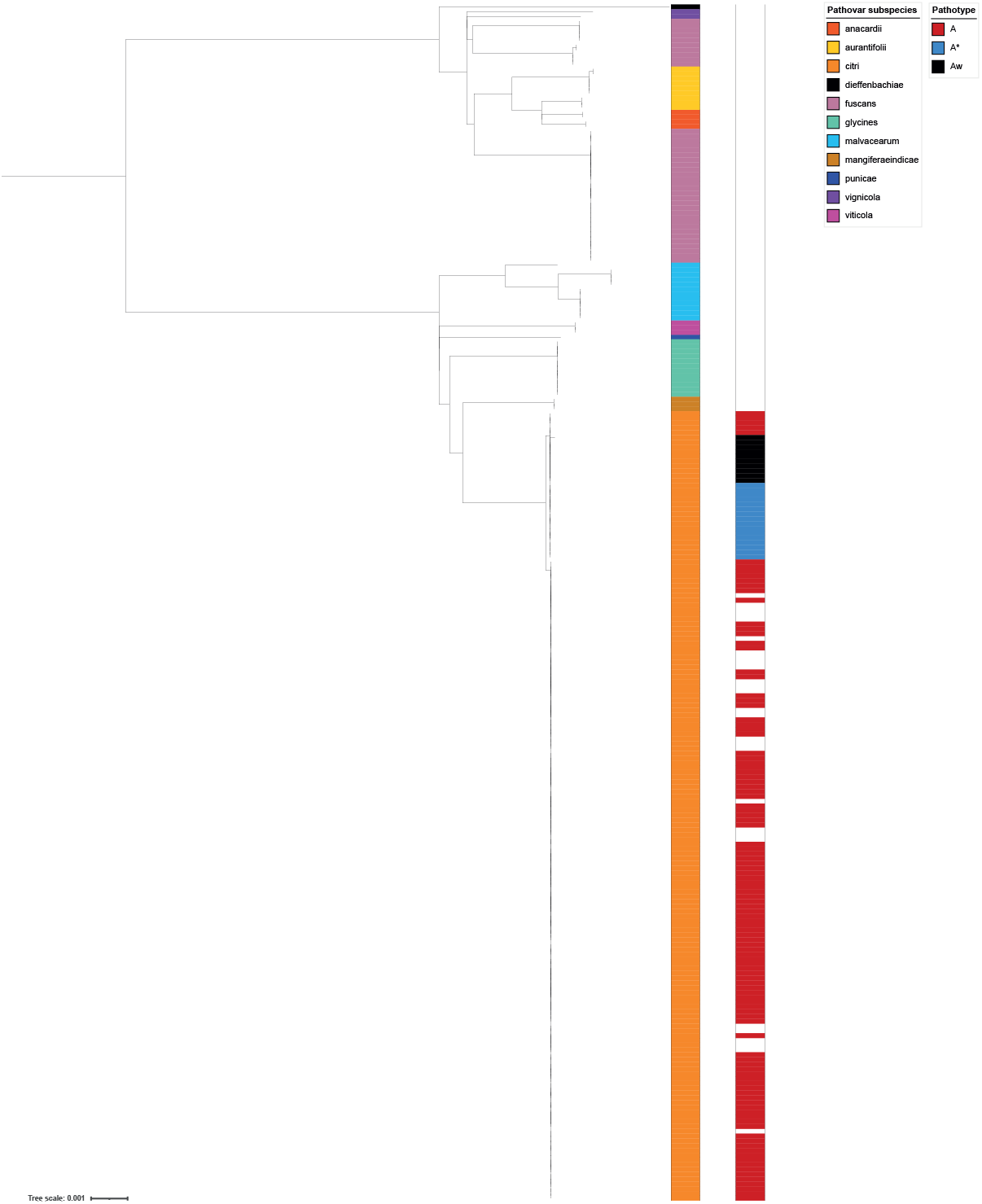
Neighbour-joining tree based upon 250 concatenated core genome MLST allele sequences of *Xanthomonas citri*. Isolates are coloured according to their original pathovar subspecies designations and XCC pathotype. Phylogeny was generated using the iTOL^(19)^ plugin at the PubMLST website (Pubmlst.org).

**Figure 2.**
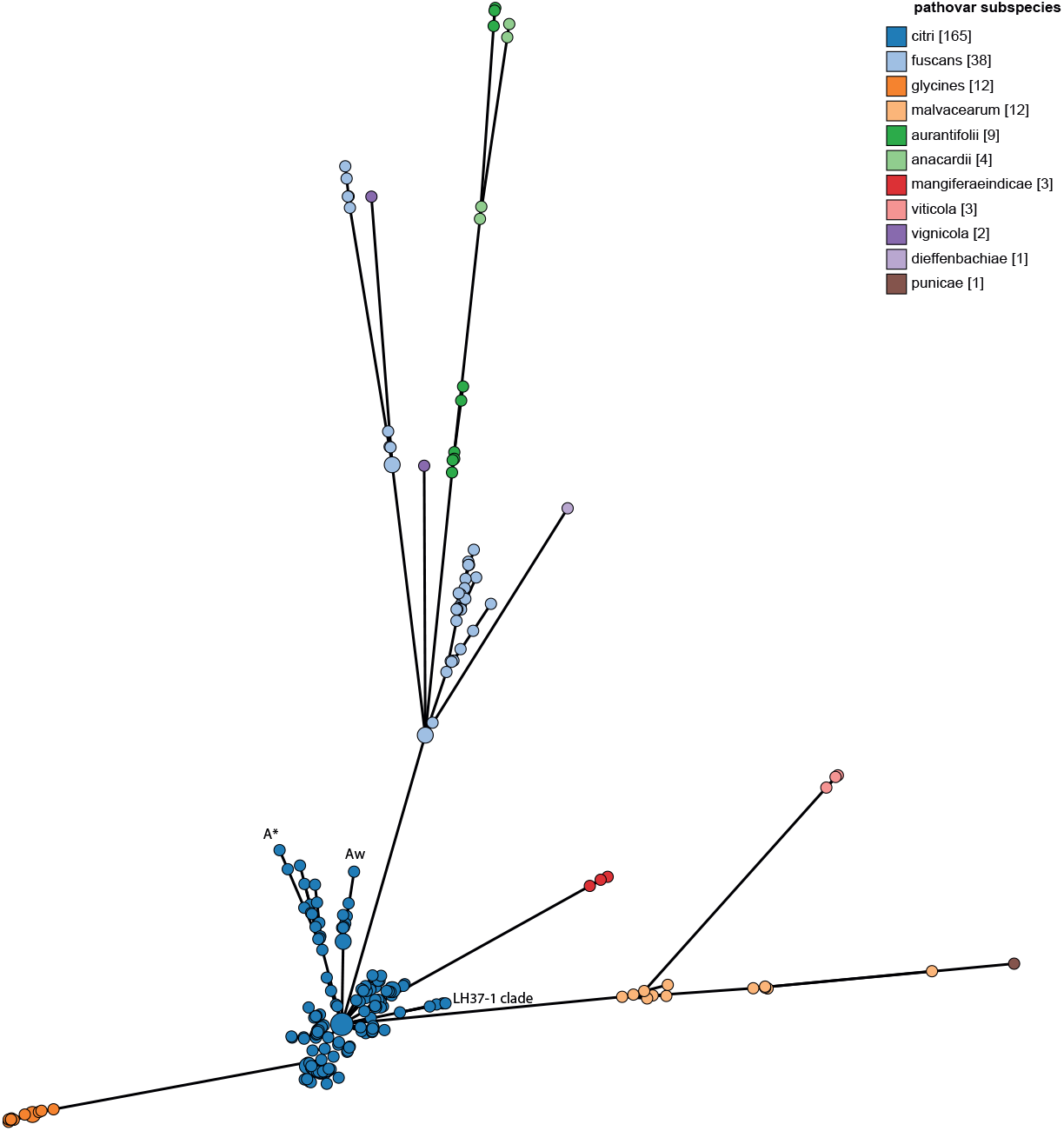
Minimum-spanning tree based upon 250 core-genome allelic profiles of *Xanthomonas citri*. Isolates / groups are coloured according to pathovar subspecies. Pathotype A*, Aw and the divergent pathotype A clade containing five genomes including that of isolate LH37-1 are shown. Phylogeny was generated using the Grapetree^(18)^ plugin at the PubMLST website (Pubmlst.org).

The time taken to generate phylogenies based on core-genome SNPs using kSNP3 and MEGA 11 (36h), concatenated allelic sequences using iTOL (15 min) and MLST allele data using Grapetree (4 min) varied considerably. All tests were run on a 2020 3.6 GHz 10-Core Intel Core i9 iMac with 16GB RAM.

### XCC Pathotype phylogenies

The genomes of pathotype A strains of pv. *citri* represented the largest group in this study and these isolate genomes resolve as a discrete group in a NJ phylogeny based upon concatenated cgMLST allele data (Figures 1 and 2) with the exception of five isolates whose genomes were listed in nucleotide submission information as being pathotype A. These five isolates correspond to a separate lineage of pathotype A isolates examined in a 2015 study by Gordon *et al*(6) that were isolated from grapefruit, key lime and citrus spp. Pathotype A* and A^w^ isolate genomes also form distinct clades using all three tree building methods (Figure 1). This can be seen in Figures 1 and 2 but is clearer in Figures 3 and 4 that include only XCC isolate genomes.

**Figure 3.**
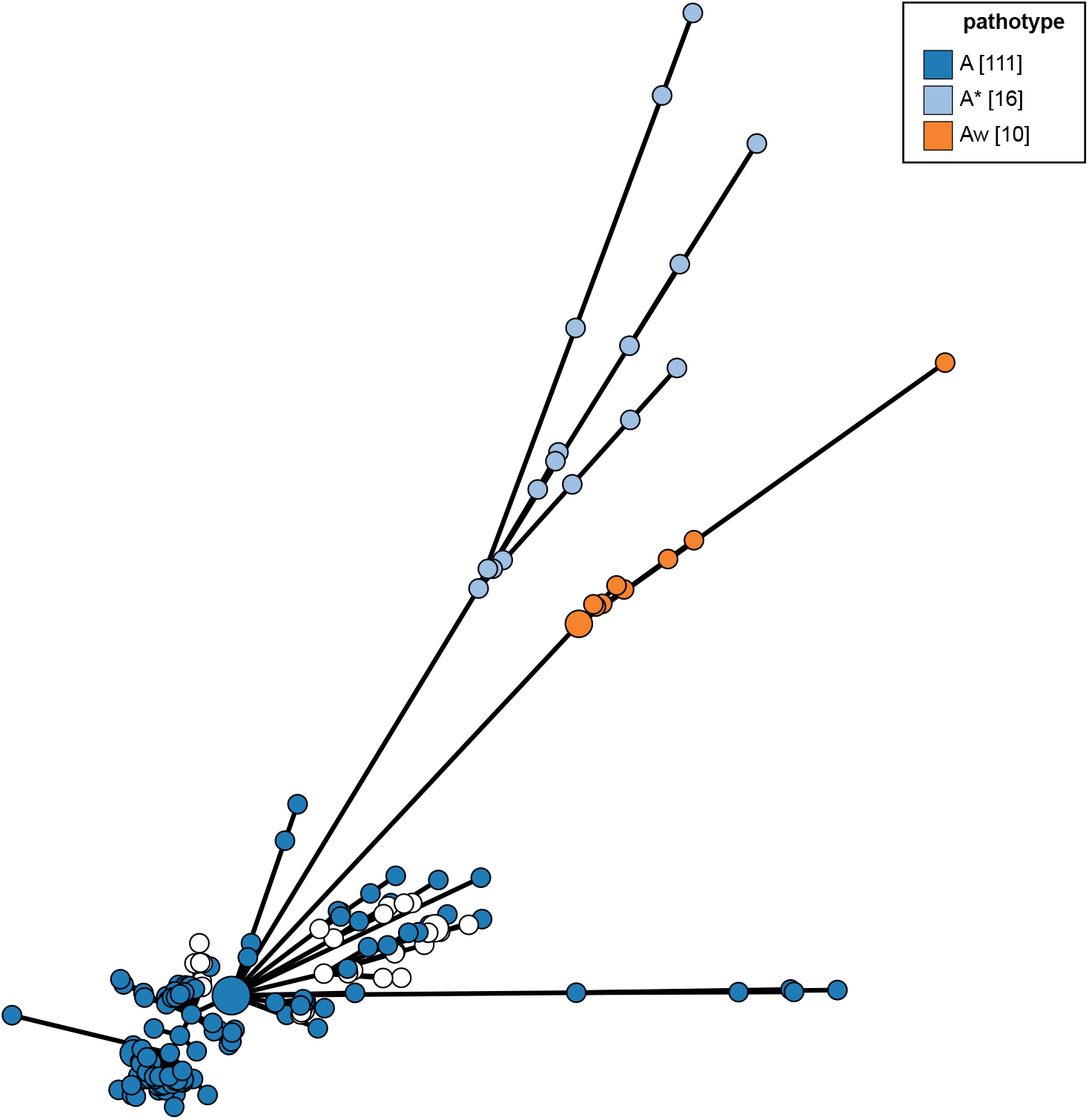
Minimum-spanning tree based upon 127 *Xanthomonas citri* pv. *citri* core-genome allelic profiles and coloured according to pathotype. Phylogeny was generated using the Grapetree^(18)^ plugin at the PubMLST website (Pubmlst.org).

## Discussion

We have developed a cgMLST scheme for the study of *Xanthomonas citri*. Through its implementation at pubmlst.org the scheme can be used to rapidly and robustly infer pathovar (or subspecies) and, in the case of XCC, pathotype designation(s) based on genetic similarity to uploaded and curated genomic sequence data with associated metadata. The database currently contains 250 isolate genomes and associated metadata including 100 novel XCC isolates sequenced in this study.

A neighbour-joining phylogeny based upon concatenated allele sequences generated using the iTOL plugin at pubmlst.org had very similar structure to a core genome SNP based NJ tree but was generated within several minutes compared to the >36h time required to generate a SNP based phylogeny that depended upon multiple whole genome alignment. This is important because, as the number of sequenced *X. citri* genomes deposited in public databases increases, the greater the computational resource required to generate SNP phylogenies *de novo*.

We used Grapetree, implemented at pubmlst.org to generate and display minimum-spanning trees of study data and this plugin can quickly and clearly display phylogenies coloured according to metadata such as country of origin, date, host species, pathovar, and XCC pathotype. Phylogenies generated using each of the methods used here - SNP based NJ, and iTOL and Grapetree phylogenies based upon concatenated cgMLST loci sequences, were largely congruent with very similar groupings. However Grapetree and its ability to easily and quickly display very large genomic data sets, such as those present in Enterobase, is eminently scalable as datasets grow, unlike core SNP based methods that are more computationally intensive and time-consuming.

The cgMLST scheme implemented at Pubmlst.org for *X. citri* will, we hope, be an increasingly useful tool for the study of the epidemiology and evolution of the major cause of citrus canker caused by *X. citri* subsp. *citri* but should also be of benefit for the study of other plant pathogenic *X. citri* subspecies and pathovars included in this study as well as those not yet included in the database.

## Financial Disclosure Statement

This work was funded by BBSRC / Newton Fund (https://www.ukri.org/councils/bbsrc/ https://www.newton-gcrf.org/newton-fund/) awards BB/R022720/1 and BB/S018891/1 to ME, CONFAP (https://confap.org.br/) research mobility award 2019/05497-7 to ME and FAPESP awards 2017/50454-9 and 2018/21164-5 to HF.

## Competing interests

None of the authors have any financial, personal, or professional interests that could be construed to have influenced the work.

## Figure Legends

**Supplementary Figure 1.**
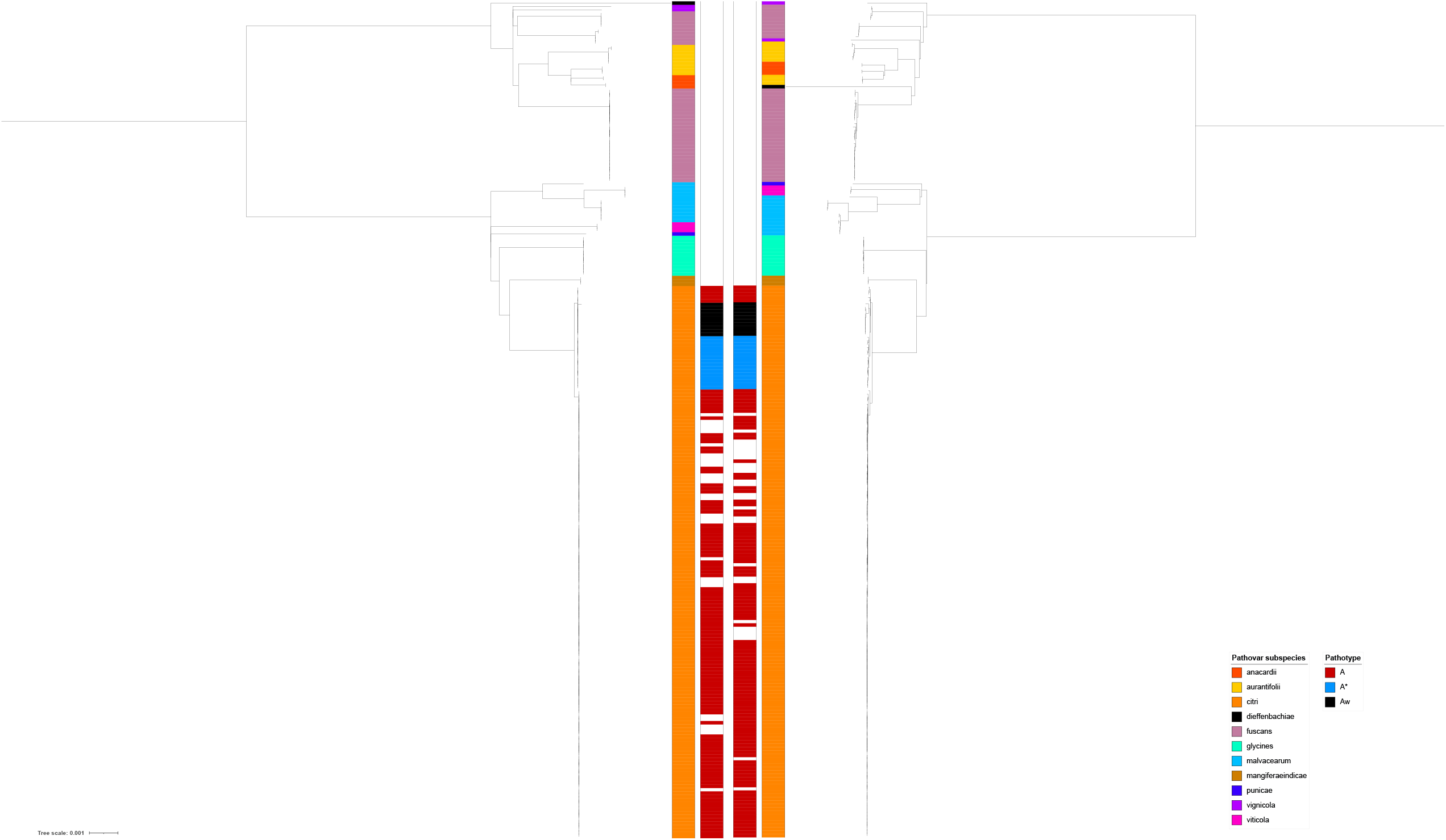
Comparison between neighbour-joining trees of 250 *Xanthomonas citri* isolate genomes generated (left) using concatenated MLST allele sequences and (right) core-genome SNPs coloured according to subspecies and *Xanthomonas citri* pv. pathotype.

